# The evolution of contact prediction: Evidence that contact selection in statistical contact prediction is changing

**DOI:** 10.1101/660191

**Authors:** Mark Chonofsky, Saulo H. P. de Oliveira, Konrad Krawczyk, Charlotte M. Deane

## Abstract

Over the last few years, the field of protein structure prediction has been transformed by increasingly-accurate contact prediction software. These methods are based on the detection of coevolutionary relationships between residues from multiple sequence alignments. However, despite speculation, there is little evidence of a link between contact prediction and the physico-chemical interactions which drive amino-acid coevolution. Furthermore, existing protocols predict only a fraction of all protein contacts and it is not clear why some contacts are favoured over others.

Using a dataset of 863 protein domains, we assessed the physico-chemical interactions of contacts predicted by CCMpred, MetaPSICOV, and DNCON2, as examples of direct coupling analysis, meta-prediction, and deep learning, respectively. To further investigate what sets these predicted contacts apart, we considered correctly-predicted contacts and compared their properties against the protein contacts that were not predicted.

We found that predicted contacts tend to form more bonds than non-predicted contacts, which suggests these contacts may be more important. Comparing the contacts predicted by each method, we found that metaPSICOV and DNCON2 favour accuracy whereas CCMPred detects contacts with more bonds. This suggests that the push for higher accuracy may lead to a loss of physico-chemically important contacts.

These results underscore the connection between protein physico-chemistry and the coevolutionary couplings that can be derived from multiple sequence alignments. This relationship is likely to be relevant to protein structure prediction and functional analysis of protein structure and may be key to understanding their utility for different problems in structural biology.

**Author summary:** Accurate contact prediction has allowed scientists to predict protein structures with unprecedented levels of accuracy. The success of contact prediction methods, which are based on inferring correlations between amino acids in protein multiple sequence alignments, has prompted a great deal of work to improve the quality of contact prediction, leading to the development of several different methods for detecting amino acids in proximity.

In this paper, we investigate the properties of these contact prediction methods. We find that contacts which are predicted differ from the other contacts in the protein, in particular they have more physico-chemical bonds, and the predicted contacts are more strongly conserved than other contacts across protein families. We also compared the properties of different contact prediction methods and found that the characteristics of the predicted sets depend on the prediction method used.

Our results point to a link between physico-chemical bonding interactions and the evolutionary history of proteins, a connection which is reflected in their amino acid sequences.

## Introduction

The development of advanced methods to detect correlation between sites in large multiple sequence alignments has increased the accuracy of protein contact prediction. The predicted contacts output by these methods have resulted in improvements in many areas of structural biology, including template-free protein structure prediction [1, 2]. Machine learning-assisted contact prediction methods, such as AlphaFold, have recently demonstrated unprecedented ability to accurately predict protein structures at the level of topology or better [3].

These contact prediction methods are based on the idea of coevolution between residues in the protein structure. If a protein is to keep its folded shape when a residue mutates, at least one of the residues with which it is in contact is likely to undergo a compensatory mutation. For example, a mutation which removes one cysteine in a disulfide bond might be compensated by a mutation of the remaining cysteine in order to preserve a bonding interaction between those two sites in the protein. Sites where such compensatory mutations occur frequently can be identified by statistical techniques from multiple sequence alignments. For these techniques to be successful, it is necessary that the multiple sequence alignments contain sufficient levels of sequence diversity to reveal these correlations.

Early contact prediction methods used mutual information between alignment columns to infer contacts. Even with a number of corrections, particularly including the average product correction [4] for phylogenetic and entropic noise, these methods (such as MIp [4], MIc and aMIc [5], and ZNMI [6]) were unable to accurately infer protein contacts (*i.e.*, residues that share spatial proximity, typically those with C_β_ less than 8 Å apart). Gomes *et al.* (2012) found less than 30% precision at 20% recall for any of the available mutual information-based methods. The low precision of these methods was due in part to their inability to identify contacts within a larger number of transitive correlations.

Direct coupling analysis (DCA) [1, 7, 8] overcame some of the weaknesses of MI methods by correcting for the effect of transitive couplings between residues. Methods such as CCMpred [9], Freecontact [10], EVFold [11], GREMLIN [12], and PSICOV [1] all use variations of this methodology. DCA-based contact predictors reached accuracies approaching 50% for the top *L/*5 contacts where *L* is the length of the protein [1]. Despite higher accuracy, these methods still obtains a low recall, and it remains unclear why certain contacts are not predicted. Hockenberry *et al.* [13] have suggested that DCA methods detect side-chain interactions.

In an effort to further increase accuracy and recall, the next development in protein contact prediction was the introduction of meta-predictors, which combined the output of different contact predictors to create aggregate predictions (*e.g.* MetaPSICOV [14] and PConsC [15]). MetaPSICOV outperforms its constituent predictors (CCMpred, DCA, and PSICOV) by 10% precision or more, as assessed on the top *L* contacts [14]. Although these methods increase the number of correctly predicted contacts, they also predict a set of contacts which is different from the sets that their constituent predictors predict, for example, by removing contacts that are predicted with low confidence or by only one constituent predictor, or by ‘filling in’ contacts from secondary structures [14].

The most recent developments have been the application of deep learning approaches to contact prediction. DNCON2 [16] and RaptorX [17] are currently the only published examples of deep learning based contact predictors. (CASP13 featured numerous examples of this class of approach, but these programmes have not yet been released to the community.) Neither RaptorX nor DNCON2 operates directly on the multiple sequence alignment, instead using features derived from statistical coupling inference methods and sequence property predictions, such as predicted secondary structure and predicted solvation. DNCON2 outperforms MetaPSICOV and RaptorX on the CASP10, CASP11, and CASP12 datasets [16], achieving a precision of 53.4% on the CASP12 dataset, compared with 42.9% and 46.3%, respectively, for MetaPSICOV and RaptorX, for the top *L/*5 predictions of long-range contacts. These methods treat contact prediction as a problem in computer vision, enabling the application of higher-order structures to the data, and resulting in a set of correctly-predicted contacts that is again larger than those predicted by DCA or meta-prediction methods. This larger set must again contain different contacts from those identified by DCA or meta-prediction.

Contact prediction methods have been used to approach many bioinformatics problems, from protein structure prediction to inference of functional interactions, but little work has been done to understand the nature of the contacts that they predict. Given that these methods were all initially based on identifying co-evolving sites, it could be expected that the contacts that they predict relate to specific types of interactions. It is also likely that there are differences between contacts predicted by different methods. While more modern prediction methods may improve the accuracy of the predictions, as they move further from attempting to extract coevolutionary signal, the physico-chemical nature of the sets of predicted contacts may change. Direct coupling methods identify contacts that exhibit strong statistical coevolutionary signal, and may therefore identify contacts that have particular evolutionary significance. The effect of adding other information to these predictions through deep learning is not known. These differences might be key in understanding their utility for different problems.

In this paper, we investigate the nature of the predicted contacts from different contact prediction methods. We compare aMIc, CCMpred, MetaPSICOV, and DNCON2 as examples of the different types of contact predictors currently available, and we assess the differences between true contacts predicted by the methods and random true contacts in protein structures. We classify the bonds which are formed between residues in our sets of contacts, and we show differences in the number and kind of physico-chemical bonding interactions between different methods, and between predicted contacts and random contacts. We show commonalities between machine-learning based methods (MetaPSICOV and DNCON2) and direct coupling analysis. Further, we find differences in the extent to which bonds are conserved between different sets of contact predictions and between contact predictions and the set of all contacts.

## Materials and methods

### Structural domain set

From the 13,760 domains in ASTRAL (06.02.2016 build at 40% sequence identity cut-off) [18–21], we selected a single exemplar domain for each CATH [22] homologous superfamily, giving 2,086 protein domains. For each protein domain, we assembled a Multiple Sequence Alignment (MSA) and predicted contacts for that alignment. (See below for more details.)

### Multiple sequence alignment generation

For each domain, we generated an MSA using HHBlits 3.0.0 (15-03-2015, default options except `-n 3, -maxfilt 500000, -id 99, -cov 0.90`) with the Uniprot20 database (2016.02) [23]. In order to ensure alignments of sufficient quality for use in contact prediction, we removed MSAs which had *N*_*f*_ < 32 [24].

### Contacts

Contacts are defined as residue pairs where the distance between C_β_ atoms (C_α_ for glycine) is less than 8Å. While this cut-off is arbitrary, it is in accordance with convention in the field, and in particular it is the cut-off with which DNCON2 and MetaPSICOV were trained [14, 16]. We consider only those contacts which are separated by five or more residues.

### Contact prediction

We used our MSAs as input to four contact prediction methods: aMIc [5], CCMpred [9], MetaPSICOV version 1 [14], and DNCON2 [16]. For each of these prediction methods, we used default parameters except in the following ways. For aMIc, we used a pseudocount value of 0.05 in pairwise residue counts so that the marginal contributions of the pseudocounts for each residue was 1. We also modified the DNCON2 pipeline to use our HHBlits alignments so that all four methods had identical input. After contact prediction, we assessed contact prediction accuracy and removed cases in which any of CCMpred, MetaPSICOV, and DNCON2 had contact prediction accuracy over the top *L* contacts below 30%, where *L* is the length of the protein domain. We also removed structures where there were too few real contacts to populate the background set (see below). A full list of all 2,086 cases and their alignment and contact prediction statistics are given in S1 Table.

### Physico-chemical interactions

We used ARPEGGIO [25] to identify the types of physico-chemical interactions between amino acids in the three-dimensional protein structures of our domains. ARPEGGIO uses molecular geometry to classify physico-chemical interactions into 13 Structural Interaction Fingerprints (SIFts) [26]. The most common interaction types by overall count were `hydrophobic; polar, hydrogen bond, and weak polar and weak hydrogen bond;` and `vdw` (van der Waals). We also observed `carbonyl, aromatic, ionic`, and `covalent` interactions. We did not count the `proximal` category because it is a *d* ≤ 5*Å* distance bin, overlapping substantially with other interaction types without implying a specific physico-chemical interation. A full list of physico-chemical interaction types is given in SI Table S2 Table. We call these attractive physico-chemical interactions “bonds” because they represent attractive physical interactions between atoms. While some (i.e., disulfide bonds) are covalent, most are not.

### Structural alignment

Protein-protein structural alignments were carried out with CATH-SSAP [22], since we used CATH homologous superfamilies in structural classification.

### Secondary structure classification

STRIDE [27] was used to assign contacts to secondary structures. We classified contacts into four categories: Loop-Loop (contacts formed between residues in loops), SS-Loop (contacts formed between a residue in a loop and a residue in a secondary structure elements), within-SS (contacts formed between residues within one secondary structure element), and between-SS (contacts formed between residues within two different secondary structure elements). We classified contacts as within-SS by considering runs of consecutive *α* or *β* residues. If two contacting residues *A* and *B* were situated in runs *R*_*A*_ and *R*_*B*_ of the same secondary structure type, we classified the contact (*A, B*) as within-SS if there was a main-chain hydrogen bond between any of the residues in *R*_*A*_ and *R*_*B*_, or if *A* and *B* were situated in the same run. We also allowed transitive effects: if a third residue *C* were located in a run *R*_*C*_ that had a main-chain hydrogen bond with *R*_*B*_, the contact (*A, C*) would have been classified as within-SS.

### Effective isolated contacts

To assess the distribution of contacts, we sought the largest set of contacts which could be considered isolated. Specifically, we considered a contact (*A*_1_, *A*_2_) between amino acid *A*_1_ and amino acid *A*_2_ to be isolated if there was no predicted contact (*B*_1_, *B*_2_) such that |*A*_1_ − *B*_1_| ≤ 1 and |*A*_2_ − *B*_2_| ≤ 1. We constructed an undirected graph on predicted contacts, with contacts corresponding to vertices and edges between contacts *A* and *B* iff |*A*_1_ − *B*_1_| ≤ 1 or |*A*_2_ − *B*_2_| ≤ 1. We then found a minimal vertex cover on this graph using a 2-approximation algorithm [28], *i.e.*, we identified the minimal set *C* of contacts such that *C* was adjacent to every contact not in *C*. The number of effective isolated contacts was the number of contacts not present in the vertex cover. We computed the vertex cover for all correct contacts inferred by any method.

### Adjusted probabilities

We computed the probability that a contact of a given bond type was predicted by a given prediction method. In order to account for different sizes of contact sets from different prediction methods, we adjusted these probabilities by a factor equal to the ratio of the length *L* of the protein to the number of correct contacts in the set under consideration *i.e.*,

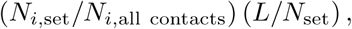

for a bond type *i* and the number *N*_set_ of contacts in the predicted set for a contact prediction method. These probabilities are scaled to compensate for the effect of predicted sets of different sizes due to different contact prediction accuracies. These adjusted probabilities were averaged over the 863 cases.

### Terminology

#### Predicted set

Of the top *L* predicted contacts for a given protein structure, the predicted set is the set of residue pairs which are in contact in that protein structure (true contacts), where *L* is the length in residues of the structure. Therefore, the size of the predicted set is at most *L*.

#### Background set

A randomly-selected set of residue pairs which are in contact in a given protein structure. For each protein structure, we select the same number of contacts for the background set as are in the predicted set, and we exclude residues which are in the predicted set. For most analyses, we use 20 randomly-selected background sets for each structure.

## Results

### Trends in contact prediction accuracy

We predicted contacts on 1,030 protein domains which had high-quality alignments using four contact prediction methods (aMIc, CCMpred, MetaPSICOV, and DNCON2) (see Figure 1). Figure 2 shows the accuracy achieved over the top *L* contacts, where *L* is the length of the protein. As expected, aMIc (the mutual information method) performed worst (average accuracy of 15%). The best-performing method was DNCON2 (average accuracy of 77%) followed by MetaPSICOV (average accuracy of 64%) and CCMpred (average accuracy of 47%). We found that alignment quality was correlated with prediction accuracy for all prediction methods (S1 Fig). Since the purpose of this study is to investigate the physico-chemical properties of the true predicted contacts, we did not take aMIc contact predictions forward for further analysis, as only 102 cases had top-*L* accuracy equal to 30% or higher. To fairly compare the three methods in terms of the physico-chemical properties of their predicted contacts, we used only the 863 cases for which all three methods had top-*L* prediction accuracy above 30% and sufficient contacts available in the structure to form a predicted set and a background set for our analyses.

**Fig 1.**
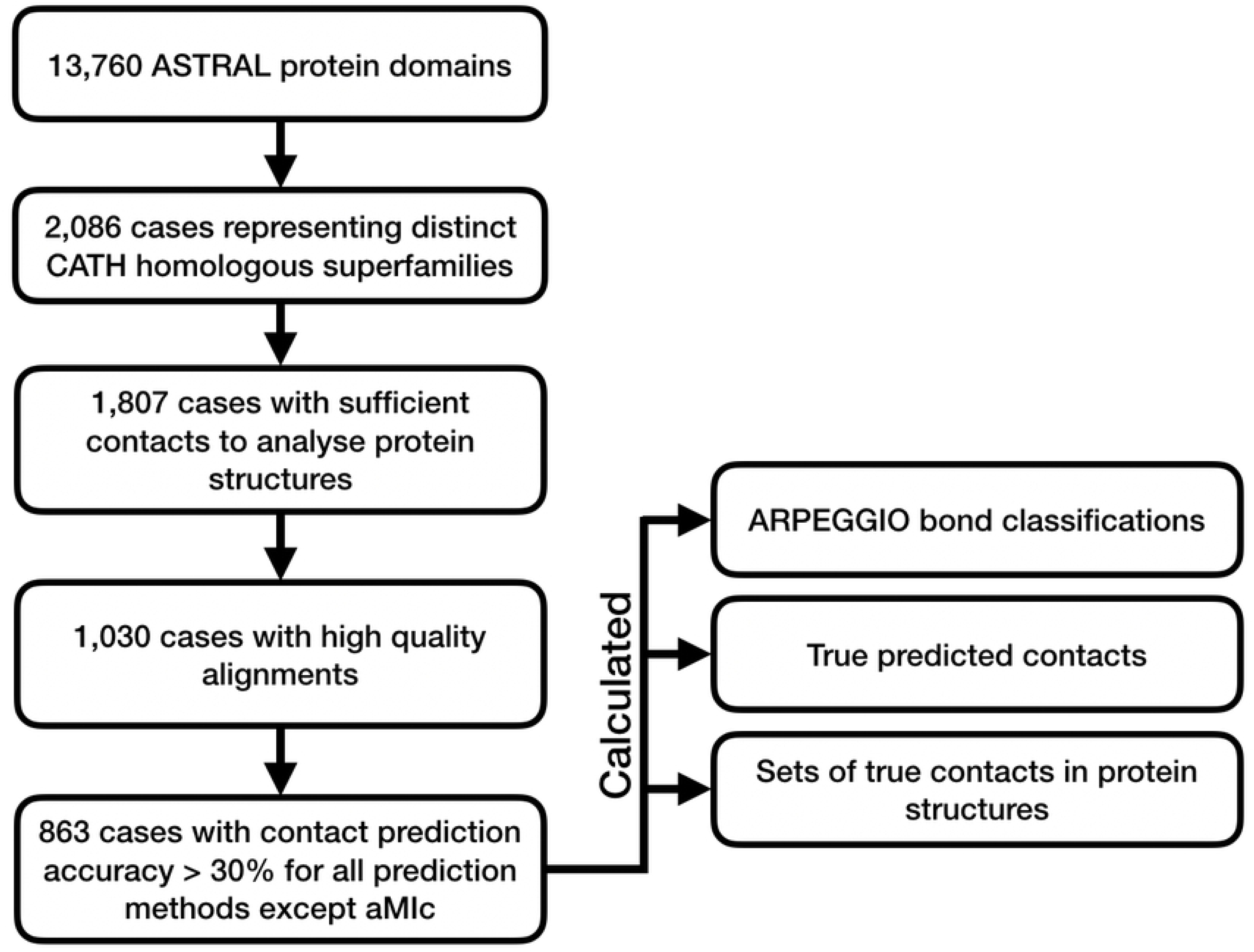
A schematic of the data processing pipeline for our analysis. As described in the main text, we filtered domains from ASTRAL to produce a set of domains with structural and functional diversity. This set of domains was used as the basis for contact prediction and categorisation of structural properties.

**Fig 2.**
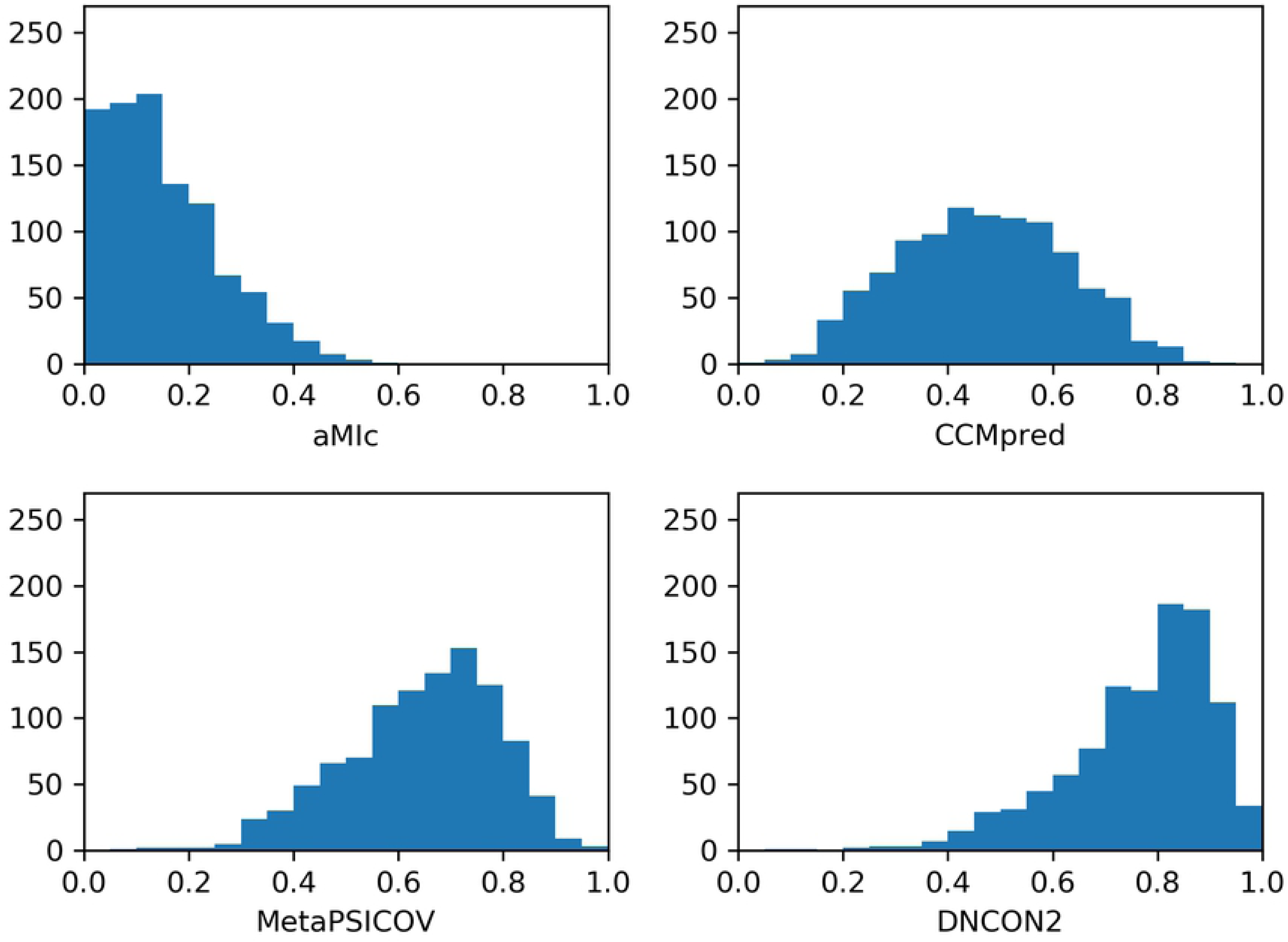
Top-*L* accuracy histograms of different contact prediction methods. Accuracy was computed with respect to the top *L* scoring predictions, where *L* is the length of the protein domain, for four prediction methods – aMIc, CCMpred, MetaPSICOV, and DNCON2 – over 1,030 protein domains. The *y* axis is the number of protein domains, and the *x* axis is the top-*L* accuracy. This analysis excludes cases where effective sequences *N*_*f*_ < 32, which is known to result in poor predictions [24].

### Predicted contacts have more bonds than background contacts

Using this set of 863 cases, we compared the properties of the correct predicted contacts for each case (predicted set) to those of a randomly-selected set of residue pairs that are in contact in that protein structure and which were not in the predicted set (background set). The bonds between residue pairs in both the background and predicted sets were identified by ARPEGGIO (see Methods). Fig. 3 shows the number of bonds per contact averaged over the 863 prediction cases. For all three contact prediction methods, there are more bonds per contact for the predicted contact sets than the background contact sets. CCMpred exhibits the largest increase (58%), while MetaPSICOV has the smallest increase (47%). The bias toward selecting heavily-bonded contacts for all prediction methods suggests that physico-chemical bonds play a role in determining the coevolutionary signal in alignments. If the need to preserve existing chemical interactions drives the correlated mutations that give rise to the evolutionary signal in protein multiple sequence alignments, then it makes sense that those contacts which have the largest number of bonds generate are likely to be predicted, and that introducing other sources of contacts would result in fewer bonds per contact.

**Fig 3.**
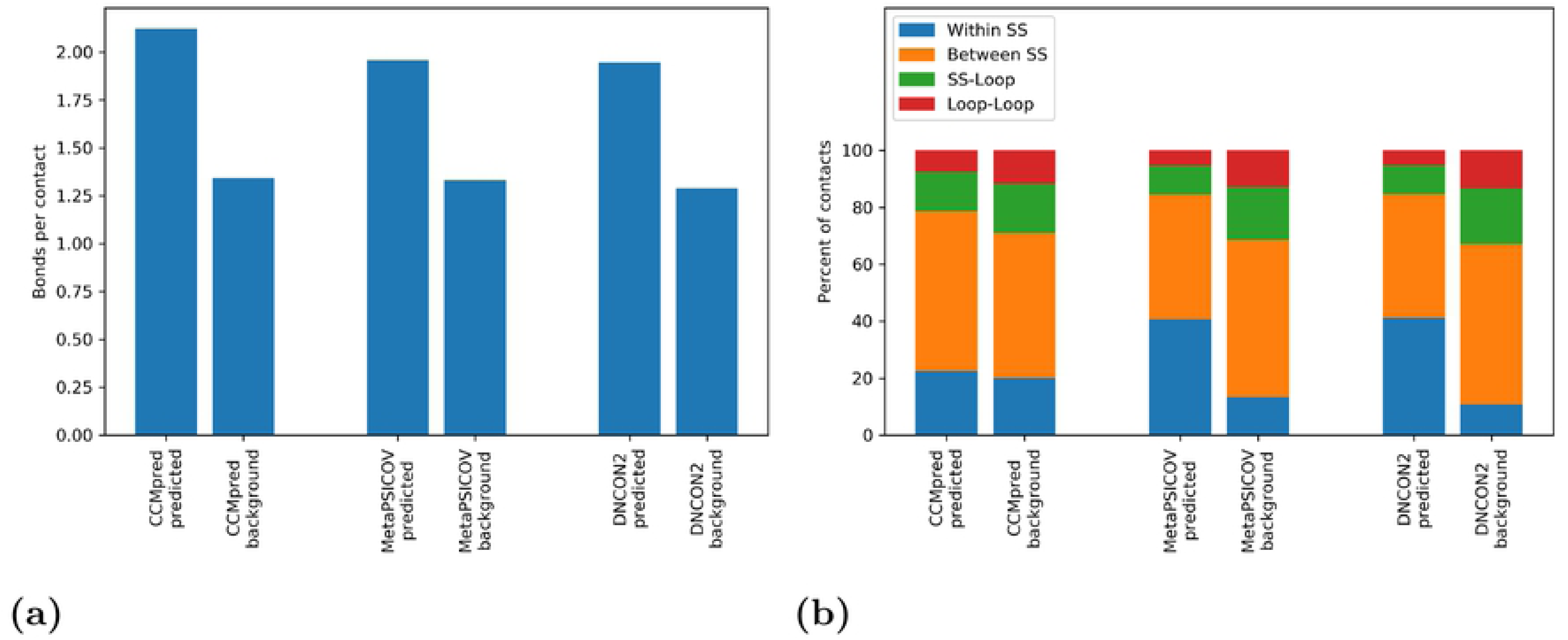
A comparison of interactions between predicted set and background set contacts. (a) shows the number of bonds per contact for the prediction methods in terms of the background and predicted sets of contacts. The figure shows the average value of bonds per contact 863 protein domains with top-*L* prediction accuracy above 0.3 for all three methods. (b) shows the difference in secondary structure composition of contacts between the predicted and background sets for different prediction methods. The average count of contacts between secondary structures, within secondary structures, between loop regions (Loop-Loop), or between loops and secondary structure (SS-Loop), is plotted.

### MetaPSICOV and DNCON2 predict almost twice as many within-secondary-structure contacts as CCMpred

To further probe the nature of this difference, we separated the counts of contacts that occurred between loops and secondary structures. Although contacts in general are disproportionately found between secondary structure elements, MetaPSICOV and DNCON2 predict almost twice as many within-secondary-structure contacts as CCMpred, despite their background sets having similar compositions (Fig. 3 (b)). These general measures of the sets of all contacts mask sharper effects of individual contact predictors because all contact predictors predict some of the same contacts. In order to more precisely identify the properties of individual contact predictors, we considered those contacts which were predicted only by particular contact predictors.

For each of the 863 protein domain cases, and restricting ourselves to the top *L* predictions, we considered separately those correct contacts that were predicted uniquely by CCMpred, DNCON2, and MetaPSICOV. We also considered those contacts that were predicted by pairs of contact predictors, and those which were predicted by all three contact prediction methods. We first considered the average number of contacts which were predicted by multiple contact predictors in terms of the ratio of the number of contacts predicted correctly to the length *L* of the protein. The largest group is the set of contacts that are predicted by all three methods (27%). DNCON2 and MetaPSICOV share an equivalently large number of contacts (also 27%) while CCMPred shares with MetaPSICOV and DNCON2 only 7% and 6% of *L*, respectively. This points to a strong link between DNCON2 and MetaPSICOV predictions. Moreover, MetaPSICOV has the lowest proportion of unique predictions (11% of its correct predictions), while DNCON2 and CCMPred have comparable proportions (24% and 22%, respectively), despite DNCON2’s higher predictive accuracy. This analysis points to differences between raw DCA-based methods and methods which incorporate information from other sources. DNCON2 and MetaPSICOV predict similar sets of contacts, while the CCMpred predicted sets tend to contain different contacts than the other two predicted sets. In light of the broader trend that CCMpred tends to predict fewer within-secondary structure contacts, and that there are similarities between the predictions of DNCON2 and MetaPSICOV that are not shared by CCMpred, we repeated earlier analyses to consider their distribution over those contacts that were predicted uniquely by one predictor, by pairs of predictors, and by all three predictors together.

First, considering the numbers of bonds per contact, we found that the contacts with the largest numbers of bonds on average were those that were predicted by all three methods. Those predicted by two or more methods also had more bonds per contact than those predicted by only one method. Of the contacts predicted by only one method, those contacts predicted only by MetaPSICOV had the lowest number of bonds per contact (1.26), while those predicted by CCMpred had the highest number of bonds per contact (1.77). Those contacts predicted by both CCMpred and DNCON2 had the highest number of bonds per contact (2.16), exceeding both sets of combinations which involved MetaPSICOV (1.85 and 1.80). As expected, in light of our findings related to secondary structures, contacts predicted by both DNCON2 and MetaPSICOV had the highest number of hydrogen bonds per contact (0.67, compared to 0.32 and 0.49 for those predicted by both CCMpred, and MetaPSICOV and DNCON2, respectively). These data confirm the idea that coevolutionary couplings are linked to the strength of the bonds between the residues that comprise them. Those contacts that are easiest to predict, in the sense that they are predicted by all three predictors, have the highest numbers of bonds per contact. This relationship is likely due to contacts with particularly strong and numerous bonds generating strong co-evolutionary signal which results in their prediction by all three methods. As noted below, there is not an unusually large proportion of within-secondary structure contacts in this group, suggesting that these predictions are not due to presence within secondary structures.

Those contacts predicted only by CCMpred have the largest number of bonds per contact of those sets from an individual contact prediction method. CCMpred uses raw co-evolutionary signal, and this signal appears to reflect the number of bonds in the contacts: the most heavily-bonded contacts are those which CCMpred successfully predicts.

We also assessed the secondary structure characteristics of the predicted contact sets. We note that the set with the highest level of contacts within a secondary structure (52%) are between DNCON2 and MetaPSICOV. The lowest level of within-secondary-structure contacts were those predicted by CCMpred alone (5%), followed by those shared between CCMpred and one of the other predictors. These data suggest that the co-evolutionary signal within secondary structures is relatively weak, presumably because these structures are harder to disrupt than supersecondary interactions. Machine-learning methods may also capitalize on the ease with which it is possible to recognise and suggest contacts within secondary structures, increasing their proportion of these types of contacts in order to increase their total accuracy.

### CCMpred contacts are distributed more widely in protein structures

We also sought to consider the distribution of contacts within protein structures. As described in Methods, we considered a contact (*A*_1_, *A*_2_) between amino acid *A*_1_ and amino acid *A*_2_ to be isolated if there was no predicted contact (*B*_1_, *B*_2_) from the set of all predicted contacts such that |*A*_1_ − *B*_1_| ≤ 1 and |*A*_2_ − *B*_2_| ≤ 1. As a measure of the distribution of the contacts throughout the protein, we used an established algorithm to remove contacts from the contact sets until all remaining contacts were isolated. We refer to the number of remaining contacts as *effective* isolated contacts. CCMpred had more effective isolated contacts than DNCON2 (0.090*L* and 0.052*L*) and both had more effective isolated contacts than MetaPSICOV (0.033*L*). Only 6% of those contacts that were predicted by both DNCON2 and MetaPSICOV were isolated, the lowest proportion of any combination of predictors or individual predictor. These data suggest that CCMpred predicts contacts which have a broader distribution within protein structures than MetaPSICOV and DNCON2. Specifically, our evidence is that DNCON2 and MetaPSICOV tend to predict blocks of contacts corresponding to complete secondary structures. CCMpred, however, tends to make more isolated predictions. These results suggest that machine learning-based predictors are learning to ‘fill in’ secondary structure contacts. Additionally, isolated predictions are more likely to be incorrect, so predictors may learn to discard ‘riskier’ isolated contacts and promote ‘safer’ contacts which are connected to other blocks of contacts. Other papers about machine learning for contact prediction have also noted that if a residue is in contact with another, then their neighboring residues are more likely to be in contact [29] and it appears that this effect is incorporated into DNCON2 and MetaPSICOV. These differences between bond numbers and between kinds of contacts among the contact predictors led us to consider whether bond types differed in similar ways.

### Types of bonding interactions differ between contact predictors

Predicted contacts have more bonds, which suggests a link between coevolutionary signal and the physical effects which bonds mediate. We sought to investigate whether this difference also manifested in a change in physico-chemical properties of the bonds that mediate contact predictions. We used the Cochran-Mantel-Haenszel procedure [30, 31] to test whether the distribution of bonding interactions in the background sets of proteins were different from the distribution of bonding interactions in the predicted set. In all cases, *p* ≪ 0.01, so we considered the differences between the predicted and background sets in further detail.

We considered the probabilities that a contact with a particular type of bond would be found in the predicted set using the adjusted probability methodology described in Methods. These probabilities are given in Table 2. (Probabilities for the background set are given in S3 Table and raw probabilities are available in S4 Table.) For each contact type, cases in which no contacts of that type were found in the protein structure were excluded from the average. A difference between contact prediction methods is evident from these data. The range of probabilities for CCMpred is larger than the range for DNCON2 or MetaPSICOV. Moreover, CCMpred has a different distribution of conditional probabilities than the other two contact prediction methods, where the figures are broadly similar. The contacts most likely to be selected in the top *L* are those which display `covalent` or `ionic` interactions. `carbonyl` interactions are the least likely to be chosen by CCMpred. These results suggest that CCMpred preferentially predicts stronger bond types, once again pointing to CCMpred contacts being more closely related to evolutionary significance.

**Table 1.**
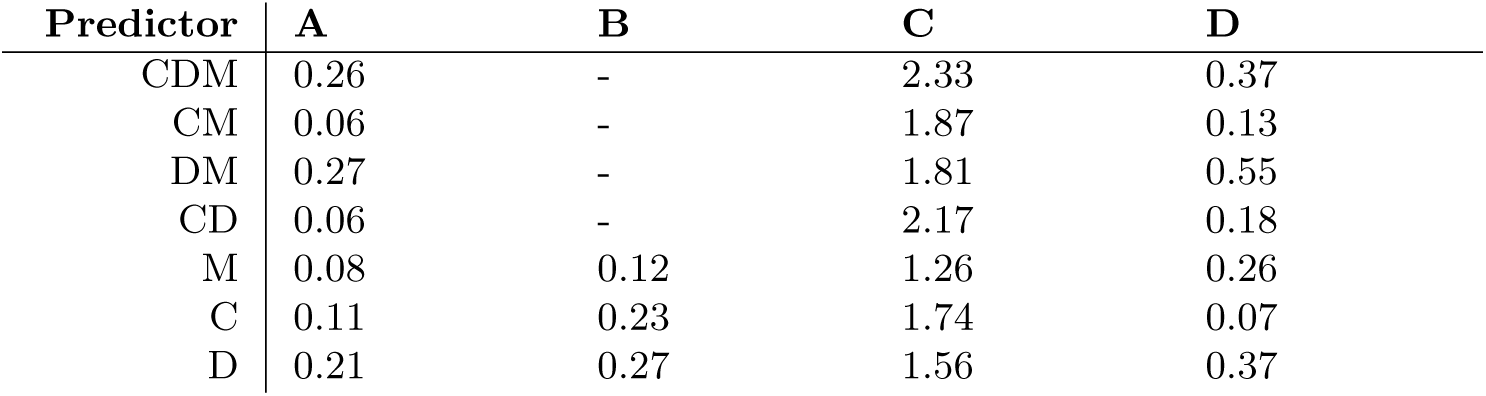
Statistics for each contact predictor. Contact prediction methods are abbreviated to their initial letters. CDM indicates those contacts predicted by CCMpred, DNCON2, and MetaPSICOV, while CM indicates those predicted only by CCMpred and MetaPSICOV, and so forth. All statistics are averages over 863 cases. **A:** Ratio of number of correct contacts to *L*, the length of the protein. **B:** Ratio of the number of contacts that are unique to each predictor to the number of the contacts predicted correctly by that contact predictor. **C:** Number of bonds per contact. **D:** Ratio of contacts within secondary structures to the number of contacts in each category, following the definition given in the main text.

**Table 2.**
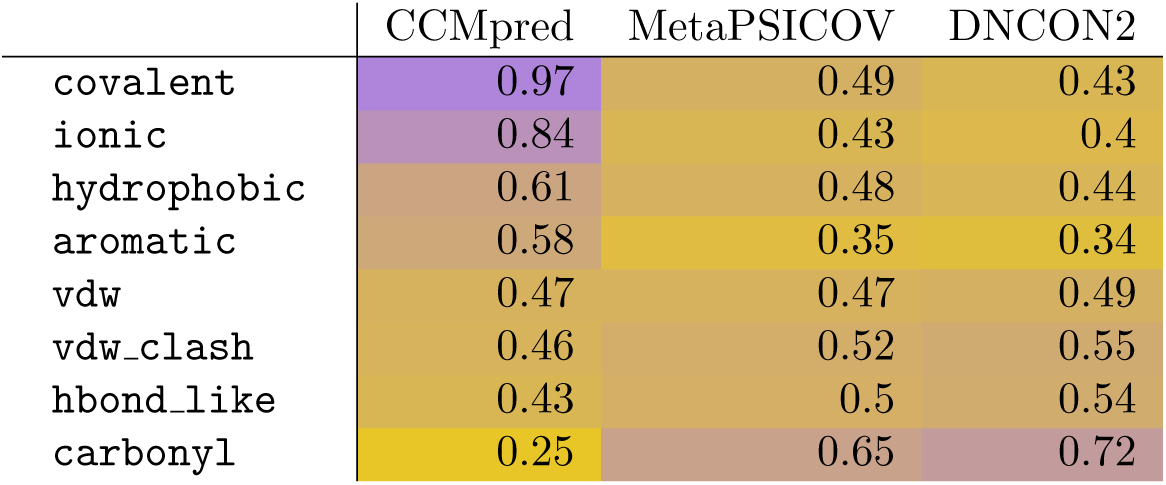
Adjusted conditional probabilities of predictions of bond types. The average adjusted probabilities that a bond of a particular type is found in the predicted set is shown in this table. These probabilities are scaled to compensate for the effect of different contact prediction accuracies as described in Methods.

### Conservation of predicted contacts

In order to further test the role of evolutionary pressure in the formation of evolutionary signal which generates these correlations, we sought to investigate whether the predicted sets were particularly highly conserved in comparison to the background sets. In order to estimate this phenomenon, we compared the extent to which the predicted set of contacts for each case *P* were present in other members of the same CATH homologous superfamily. For the CATH homologous superfamily in which *P* occurred, we filtered the homolous superfamily at a 90% sequence identity threshold and then performed structural alignment between every protein remaining in the homologous superfamily and *P*. (There were 155 CATH superfamilies which had more than one family member after filtering at 90% sequence identity.) We then recorded the proportion of the contacts in the predicted set of *P* that were also correct in the aligned family member. We performed the same process for the contacts in the background set. For all three contact prediction methods, the contacts in the predicted sets were more conserved than the background sets for more than 70% of protein-family member pairs (S5 Table). This excess was present for a range of CATH-SSAP alignment scores and grew as family members became more distant from the exemplar. Fig. 4 demonstrates how, as structural relationships become more distant, the predicted set of contacts is more strongly conserved than the background set. This effect is stronger for DNCON2 and MetaPSICOV than for CCMpred. This analysis confirms the centrality of coevolutionary constraints on our ability to predict contacts. Those contacts which are less evolutionarily important and therefore less evolutionarily conserved are more present in the background set than the predicted set. This effect is persistent over the full range of structural similarity scores within proteins. Moreover, CCMpred evinces a lower difference, which varies less as a function of alignment score than the other contact predictors. This difference may originate in CCMpred’s comparative bias against secondary structure sites, causing the predicted set to appear to be less strongly conserved than for MetaPSICOV or DNCON2.

**Fig 4.**
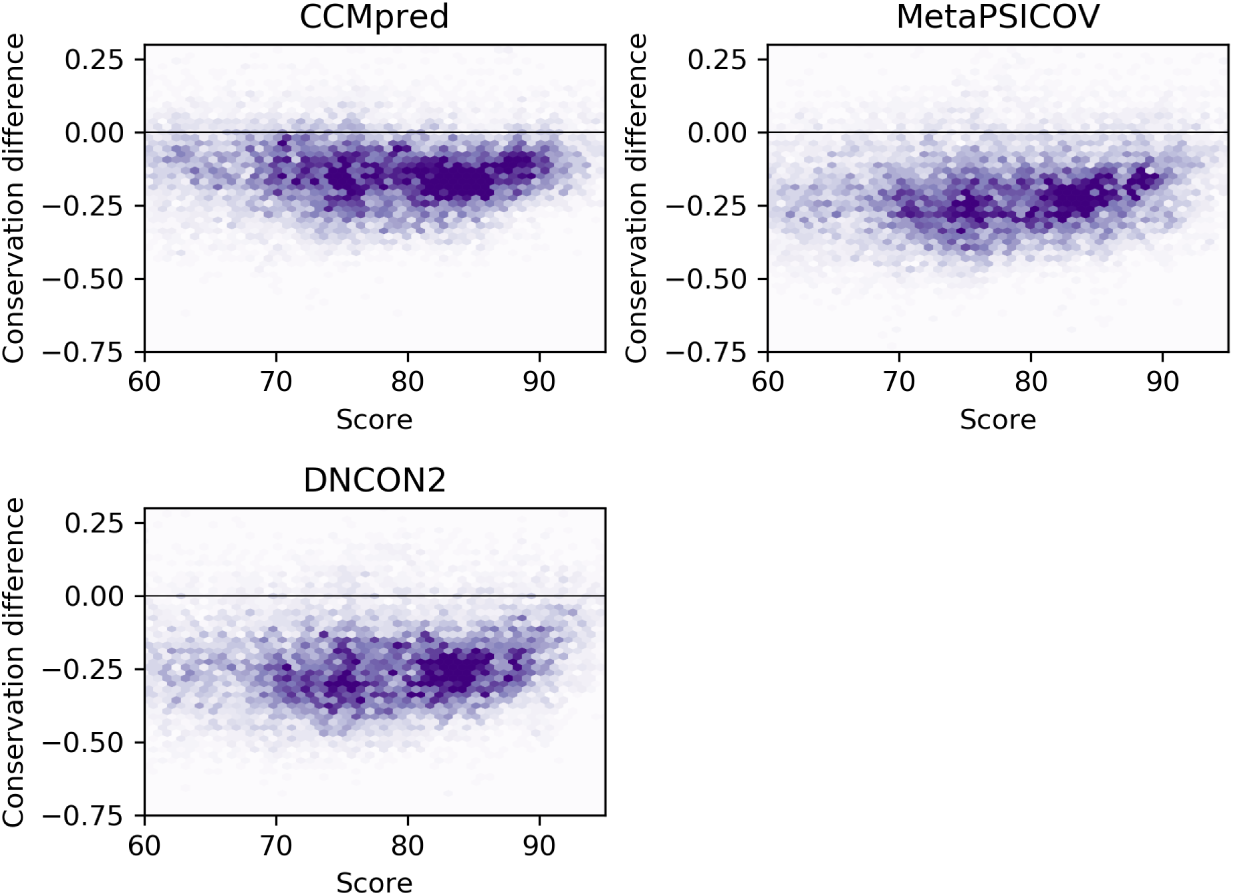
Difference in conservation between predicted set of contacts and background set for different contact predictors as a function of structural dissimilarity. SSAP structural alignment score is used as a measure of structural dissimilarity. The *y* axis is backround conservation *-* predicted conservation.

## Conclusion

Over the last ten years, contact prediction has seen remarkable gains in the accuracy of its predictions and its utility for biological applications. The field of contact prediction has been able to identify larger numbers of contacts, and our results show that this improvement has resulted in changes to the kinds of contacts predicted by state-of-the-art methods. These differences complicate the recent drive to increase prediction accuracy because not all predicted contacts may be of the same importance. In this paper, we have placed the differences between predicted and non-predicted contacts in their structural and physico-chemical context.

We found that predicted contacts and background contacts have different properties. Predicted contacts have more bonds than background contacts. For MetaPSICOV and DNCON2, more predicted contacts are within secondary structures than are background contacts. Considering those sets that are uniquely predicted by one contact predictor, these effects are heightened: the unique predictions of CCMpred have more bonds than the unique predictions of MetaPSICOV or DNCON2 and fewer within-secondary structure contacts. CCMpred contacts were more often unique to CCMpred than were MetaPSICOV or DNCON2 unique to those contact predictors. Further, CCMpred contacts were more widely distributed within the protein structures. Contact prediction methods varied in terms of the kinds of bonds that they favoured. These effects throw into relief the relationship between contact prediction and chemical bonds.

Structural constraints that are relevant to the evolutionary history of proteins, and which can be detected in multiple sequence alignments, must be mediated by some kind of physical effect. Our evidence suggests that one component of this effect are physico-chemical bonding interactions, which can be inferred from three-dimensional protein structures. These effects manifest as changes in chemical properties of contact predictions.

If contact prediction is used in the inference of structural properties, such as in the prediction of functional properties, studies of protein mechanism, or simply in structural prediction, future work must take note of the implications on contact type that their choice of prediction method entails.

The accuracy and location of predicted contacts are known to have an important effect on protein structure prediction accuracy. For this reason, a great deal of effort has been dedicated to improving the accuracy of protein contact prediction. However, our data suggest that the raw evolutionary signal of less advanced and less accurate methods may be a source of independently interesting biological information.

## Supporting information

**S1 Table. Protein family domain list.** This file contains a list of all of the protein domains considered in the analysis. It lists a collection of statistics for the alignment, including length, alignment size, effective alignment size *N*_*f*_, and contact prediction accuracy for each prediction method.

**S2 Table.**
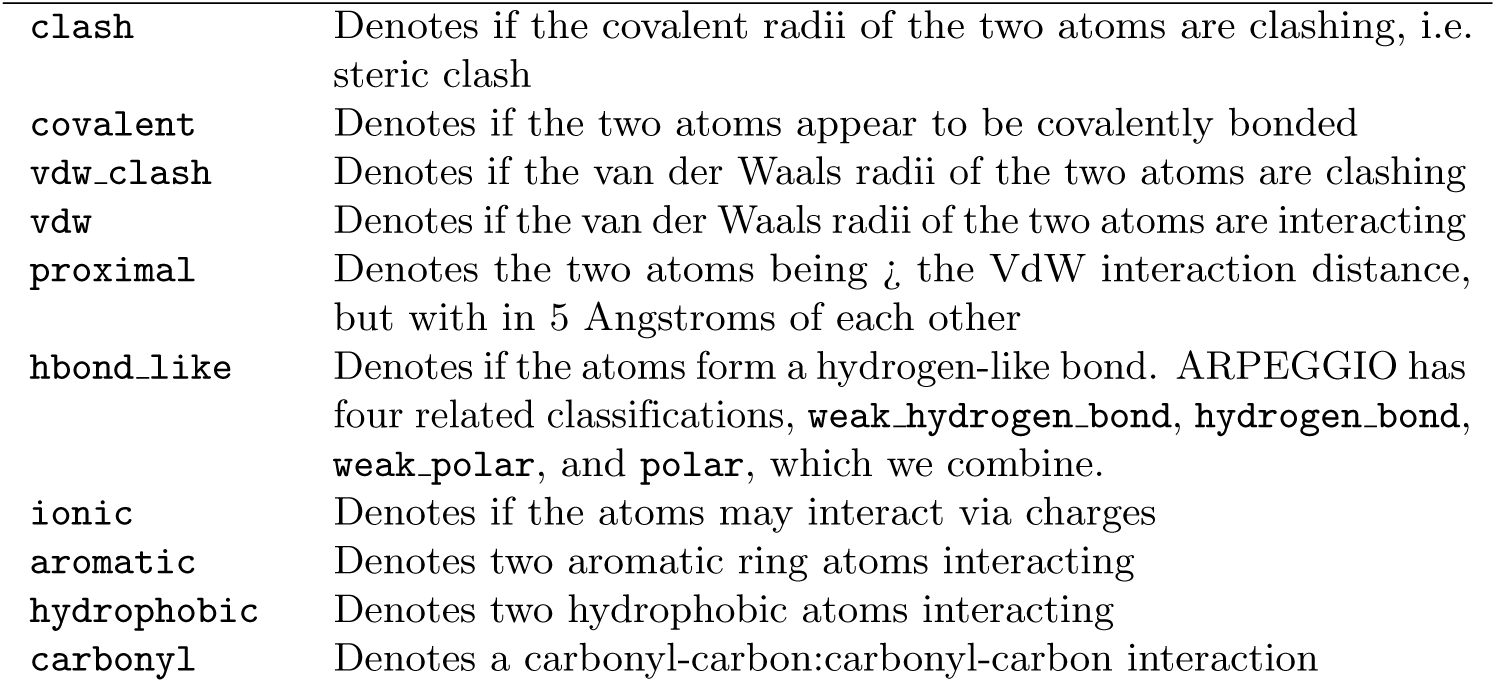
Structural Interaction Fingerprints. This table gives the SIFts identified in our collection of protein domains. SIFts are defined according to the definitions in Jubb et al. (2017) [25] except that we we combine the hydrogen bond, weak hydrogen bond, polar bond, and weak polar bond categories into one category hbond like. Identification of interacting pairs is on the basis of bond geometry and atom type. Further specification of the identification of interactions is available in [25].

**S3 Table.**
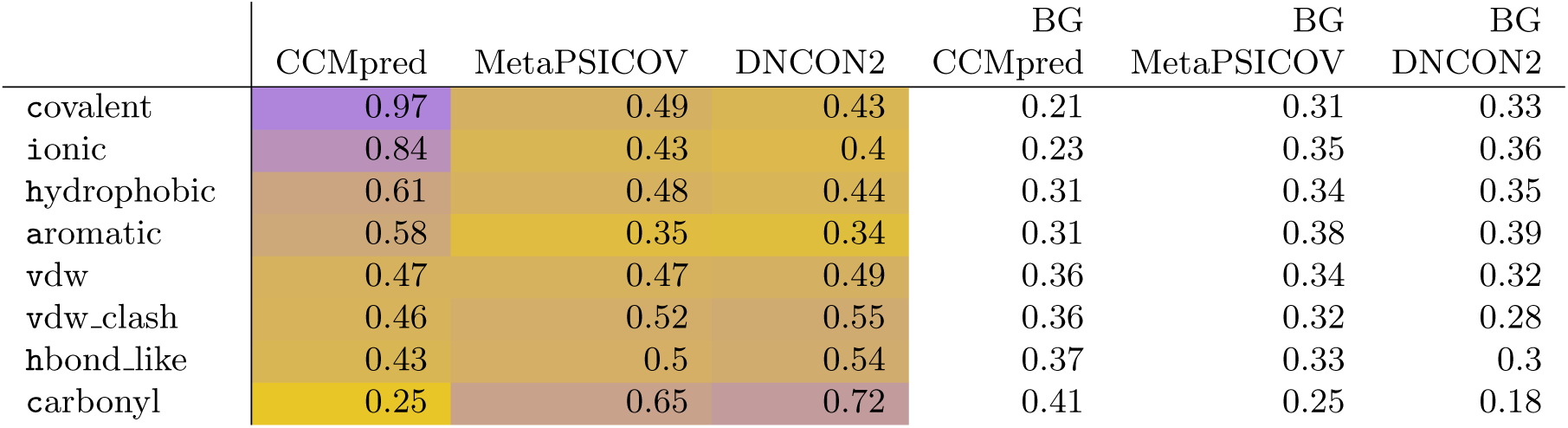
Adjusted probabilities of prediction of bond types for background and predicted sets. Table 2 gives adjusted condititional probabilities for finding a bond in the predicted set, given that it is of a certain type. In this table we give the probability of finding a bond in the background set, given that it is of a certain type. We also repeat the probabilities from Table 2 for comparison.

|  |  |  |  | BG | BG | BG |
| --- | --- | --- | --- | --- | --- | --- |
|  | CCMpred | MetaPSICOV | DNCON2 | CCMpred | MetaPSICOV | DNCON2 |
| `covalent` | 0.97 | 0.49 | 0.43 | 0.21 | 0.31 | 0.33 |
| `ionic` | 0.84 | 0.43 | 0.4 | 0.23 | 0.35 | 0.36 |
| `hydrophobic` | 0.61 | 0.48 | 0.44 | 0.31 | 0.34 | 0.35 |
| `aromatic` | 0.58 | 0.35 | 0.34 | 0.31 | 0.38 | 0.39 |
| `vdw` | 0.47 | 0.47 | 0.49 | 0.36 | 0.34 | 0.32 |
| `vdw_clash` | 0.46 | 0.52 | 0.55 | 0.36 | 0.32 | 0.28 |
| `hbond_like` | 0.43 | 0.5 | 0.54 | 0.37 | 0.33 | 0.3 |
| `carbonyl` | 0.25 | 0.65 | 0.72 | 0.41 | 0.25 | 0.18 |

**Table S4.**
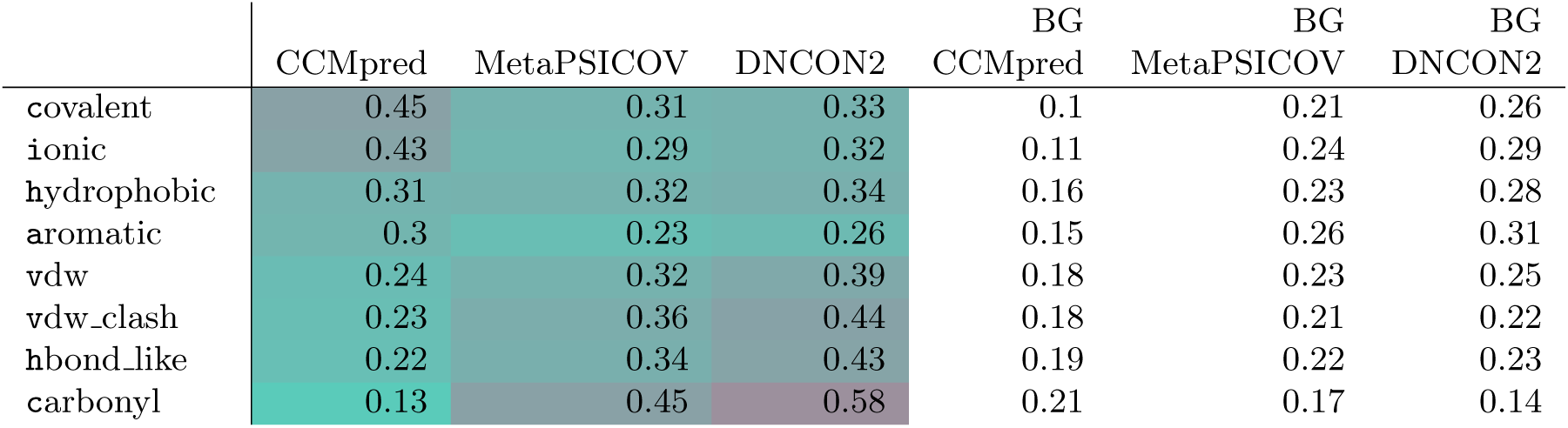
Raw probabilities of prediction of bond types for background and predicted sets. Table 2 gives adjusted condititional probabilities for finding a bond in the predicted set, given that it is of a certain type. In this table we give the probability of finding a bond in the background set. These probabilities have not been adjusted for the different average size of the predicted sets, so we would expect the probabilities for predicting each type to be lower for less accurate methods.

**Table 3. S5 Table.** As described in the main text, we compared every protein structure *i* in the CATH homologous superfamily of each our 863 prediction cases after filtering at a 90% sequence identity threshold. For each protein structure *i* in the family *J* of prediction case *i*, we computed the conservation difference 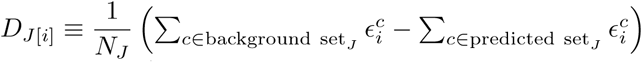, the difference between the proportion of conserved contacts in the predicted set and in the background set, where 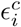 is 1 if contact *c* is a contact in structure *i* and zero otherwise. *D*_*J*[*i*]_ < 0 indicates a greater level of conservation for the predicted contacts than the background contacts. *N*_*J*_ is the number of homologues in family *J*.

| Predictor | $D \geq 0$ | $D < 0$ | Proportion $D < 0$ |
| --- | --- | --- | --- |
| CCMpred | 456 | 5906 | 92.8% |
| MetaPSICOV | 423 | 5939 | 93.3% |
| DNCON2 | 370 | 5992 | 94.2% |

**S1 Fig.**
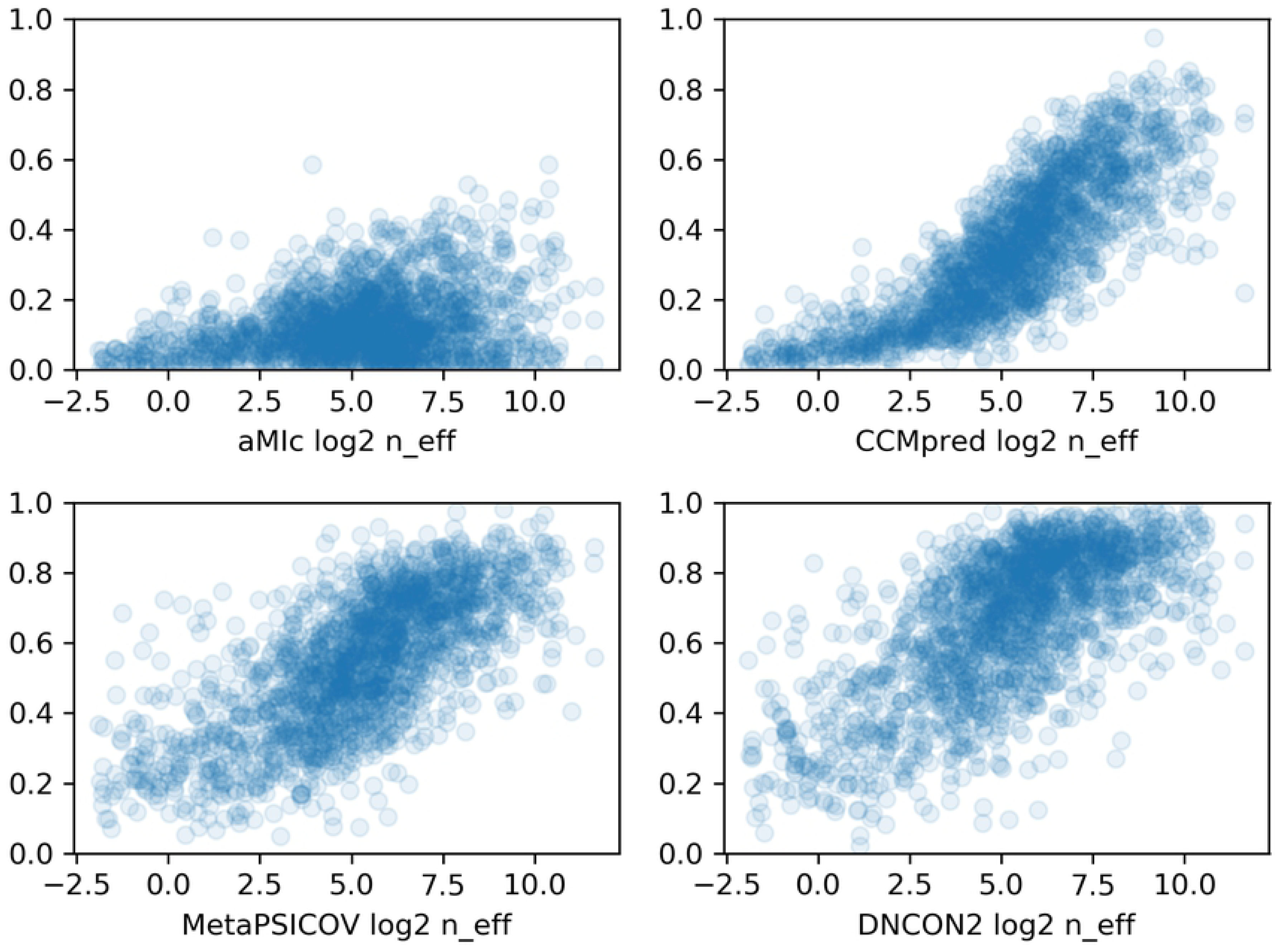
Prediction accuracy as a function of alignment quality. For each prediction method, top-L accuracy is plotted as a function of log2 *N*_*f*_ [24].

